# Comparison of polystyrene and hydrogel microcarriers for optical imaging of adherent cells

**DOI:** 10.1101/2023.10.12.559796

**Authors:** Oscar R. Benavides, Berkley P. White, Holly C. Gibbs, Roland Kaunas, Carl A. Gregory, Kristen C. Maitland, Alex J. Walsh

## Abstract

The ability to observe and monitor cell density and morphology has been imperative for assessing the health of a cell culture and for producing high quality, high yield cell cultures for decades. Microcarrier-based cultures, used for large-scale cellular expansion processes, are not compatible with traditional visualization-based methods such as widefield microscopy due to their thickness and material composition. Light-sheet tomography using label-free elastic scattering contrast from planar side illumination can achieve optical sectioning of thick samples, permitting non-invasive and non-destructive, *in toto,* three-dimensional, high-resolution visualization of cells cultured on microcarriers. Here, we assess the optical imaging capabilities of commercial polystyrene microcarriers versus custom-fabricated gelatin methacrylate (gelMA) microcarriers for non-destructive and non-invasive visualization of the entire microcarrier surface, direct cell enumeration, and sub-cellular visualization of mesenchymal stem/stromal cells (MSCs). The polystyrene microcarrier prevents visualization of cells on the distal half of the microcarrier using either fluorescence or elastic scattering contrast, whereas the gelMA microcarrier allows high fidelity visualization of cell morphology and quantification of cell density using light sheet fluorescence microscopy and tomography. The combination of optical-quality gelMA microcarriers and label-free light-sheet tomography will facilitate enhanced control of bioreactor-microcarrier cell culture processes.

## Introduction

Visualization-based monitoring of cell cultures has been a mainstay for assessing the health and quality of cell cultures for decades. The gold standard methods for quantifying cell density, for example, are based on membrane integrity and visualization of dye uptake in trypan blue dye exclusion and fluorescence live/dead assays [1], [2]. Similarly, monolayer cell culture health is routinely assessed by visualizing cells using conventional widefield microscopy techniques such as phase-contrast, brightfield, and exogenous fluorescence microscopy [3]–[8]. Furthermore, cell morphology provides information on the health, function, and fate of mesenchymal stem cells (MSCs) [9]–[15]. There is great interest in developing MSCs-based therapies due to their anti-inflammatory and immunomodulatory properties [16]–[18]. The ability to observe, quantify, and monitor cell density and morphology is imperative for assessing the health of a cell culture and for producing high quality, high yield cell cultures [19]–[22].

The anticipated widespread use of cell-based therapies, or cytotherapies, in the clinic will require manufacturing batches of trillions of cells, and one critical challenge is the efficient large-scale expansion of stem cells [23]. MSC expansion has typically been performed in 2D monolayer culture, but this is labor and reagent intensive, which limits potential for scalability. One promising strategy is to culture MSCs on spherical microcarriers maintained in suspension [24]–[26], which can produce a higher cell yield than monolayer cultures while maintaining viability, identity, and functional potential of MSCs [27]–[31]. Bioreactors reduce labor and reagent use, and have been successfully scaled up to 50 L volumes for MSC culture [23], [32]– [34]. Microcarriers can be manufactured from a wide variety of substrates with different sizes, topologies, and types of surface functionalization, such as a collagen coating or positive surface charge for enhanced attachment efficiency [35]–[39].

While there is much value in visually monitoring cytotherapeutic cells during suspension culture, standard widefield microscopy methods used to observe monolayer cultures do not readily translate to 3D microcarrier-based cultures. This is because widefield microscopes are designed to image thin monolayer cultures, whereas microcarriers can range in diameters from 100 to 300+ µm, resulting in blurred images of cells attached to the microcarrier [40], [41]. Additionally, microcarriers can be composed of materials, such as glass or polystyrene, with high refractive indices that affect the ability to visualize cells situated along the entire microcarrier surface [23], [40], [42]. Consequently, cells situated on the distal hemisphere of the microcarrier are obscured in wide-field microscopy, hence quantification of cell culture parameters like cell density and morphology are limited to imaging only one half of the microcarrier [19]. Due to the large thicknesses and relatively high refractive indices of currently available transparent microcarriers, high-resolution visualization of cells along the entire microcarrier surface requires a volumetric optical imaging technique capable of optical sectioning where a series of thin planes within the sample are sequentially imaged to reconstruct the 3D object [43]–[46].

We have previously demonstrated the advantages of gelMA microcarriers for *in toto* imaging-based quantitative monitoring of cell density and morphology of microcarrier-based cell cultures [47]. The combination of volumetric optical imaging, via planar illumination methods, and water-based hydrogel microcarriers provides the ability to rapidly and non-invasively observe the entire 3D microcarrier culture surface area, as is routinely performed for monolayer cultures [47], [48]. Light-sheet fluorescence microscopy (LSFM) is a fast and photo-gentle camera-based imaging method that utilizes fluorescence contrast for image formation. Unfortunately, LSFM requires exogenous labels for fluorescence contrast, or higher laser power for autofluorescence imaging, both of which can be toxic for live organisms [49]. Light-sheet tomography, however, presents a powerful imaging modality based on label-free elastic scattering contrast for rapid and direct visualization of individual cell morphology on microcarriers and thus enables robust non-invasive and non-destructive monitoring of cell culture critical quality attributes [47]. Since elastic scattering contrast relies on refractive-index mismatches in the sample, highly scattering microcarriers are expected to preclude the visualization of cells on their surface. The FDA has encouraged the development of process analytical technologies (PATs) for in-process monitoring of therapeutics manufacturing, and the use of label-free elastic scattering contrast would be compatible with non-destructive in- and on-line sampling of bioreactor suspension cultures, whereas fluorescence contrast would still allow monitoring of the culture but in an off-line method [50].

Most commercially available microcarriers, including those composed of polystyrene, prohibit visualization of the entire microcarrier surface due to their refractive index and opacity [19]. Here, we compare the optical imaging capabilities of commercial polystyrene microcarriers and custom-fabricated gelatin methacrylate (gelMA) microcarriers for non-destructive visualization of the entire microcarrier surface to allow direct cell enumeration and sub-cellular visualization of MSCs. Mie scattering simulations of polystyrene and gelMA microcarriers were performed to describe the angular scattering intensity distributions. The optical properties of polystyrene and gelMA microcarriers were then directly compared for compatibility with light-sheet imaging for visualization of cells and subcellular features and enumeration of adherent cells from fluorescence light-sheet and elastic scattering light-sheet images.

## Methods

### gelMA microcarrier synthesis

The gelMA microcarriers were produced following a previously published protocol [46]. Briefly, gelMA was first synthesized by reacting type A porcine gelatin (Sigma) and methacrylic anhydride at 0.6 mL methacrylic anhydride per gram gelatin in PBS for 1 hour at 50 °C. The reaction was quenched with 40 °C PBS then dialyzed through a 12 to 14 kDA nitrocellulose membrane against deionized water at 40 °C for 7 days. Then, gelMA and photoinitiator lithium phenyl-2,4,6-trimethylbenzoylphosphinate were dissolved at a final concentration of 7.5% (w/v) and 10 mM, respectively, in PBS at 50 °C. A custom polydimethylsiloxane (PDMS, Sylgard 184) microfluidic device was created and bound to a 2” x 3” glass slide after 1 minute of oxygen plasma treatment and cured overnight at 85 °C. Syringe pumps (Cole Parmer) were used to flow the gelMA solution and fluorinated oil (Novec 7500) through the microfluidic device at 2 and 4 mL per hour, respectively. The gelMA and oil droplets were then exposed to 365 nm light (75 mW per cm^2^) for 50 seconds to polymerize the microcarriers. The gelMA microcarriers were collected from the residual oil solution via centrifugation at 3000*g* for 3 min on a glycerol bed. Finally, the microcarriers were recovered from the glycerol bed and washed with PBS and stored at 4 °C.

### Bone marrow-derived MSCs cultured on polystyrene microcarriers

Bone marrow-derived human MSCs (BM-hMSCs) [51] transfected with red fluorescence protein (dsRed) were expanded on 165 µm diameter polystyrene microcarriers (Pall-Solohill) in a 10 mL rotating wall vessel (RWV) bioreactor (Synthecon) and fixed with neutral buffered formalin. First, lentiviral transduction was used to make cells express dsRed (ThermoFisher, Waltham, MA). Then, BM-hMSCs were initially expanded in low-density monolayer cell culture in complete culture medium (CCM) (α-minimum essential medium, 20% (w/v) fetal bovine serum, 2 mM L-glutamine, 100 U/mL penicillin, and 100 ug/mL streptomycin) to obtain the required cell numbers [51]. Finally, collagen I-coated polystyrene microspheres and BM-hMSCs were incubated at 1000 cells/cm^2^ at 37° C for 2 hours in 10 mL CCM with orbital mixing at 30 revolutions per minute (RPM). Cells were fixed with 4% paraformaldehyde (PFA) and stored in phosphate buffered saline (PBS) at a concentration of 3 mg particles/mL PBS. Prior to imaging, fixed microcarrier-cell samples were incubated with a 5 μM DRAQ-5 DNA staining buffer at 37° C for 30 minutes with agitation, then rinsed with PBS, and embedded in 1% agarose gel for imaging.

### ihMSCs cultured on gelMA microcarriers

Passage 4 ihMSCs were first expanded in low-density monolayer cell culture in CCM to obtain the required cell numbers. As previously described elsewhere [47], the ihMSCs [52] were cultured in a rotating wall vessel (RWV) bioreactor (RCCS-8DQ bioreactor, Synthecon) at 24 RPM on custom 120 ± 6.2 µm diameter gelMA microcarriers at 1000 cells/cm^2^ [46]. Specimens were suspended in 1 mM concentration of CellTracker Green (CTG) (Thermo Scientific) for 45 minutes, then fixed with 4% PFA, and stored in PBS. Prior to imaging, samples were incubated with a 5 μM DRAQ-5 (Thermo Scientific) DNA staining buffer at 37° C for 30 minutes with agitation, then rinsed with PBS. CTG permits visualization of the cell cytoplasm and quantification of cell morphology. DRAQ-5 is used for cell nuclei visualization and direct cell enumeration.

### Light-sheet fluorescence microscopy

*In situ* fluorescence imaging of cells attached to microcarriers was performed on a Zeiss Z.1 Lightsheet microscope (LSM) using a 20X 1.0 NA (water) detection objective lens and 10X 0.2 NA illumination objective lenses (air). BM-hMSCs were illuminated with 561 nm (DsRed) and 638 nm (DRAQ-5). ihMSCs were illuminated with 488 nm (CTG) and 638 nm (DRAQ-5). The emission filters used were 505 – 545 nm (CTG), 600 – 600 nm (DsRed) and 660+ nm (DRAQ-5). A variable zoom of 1.16x was used for an effective magnification of 23.2x and the voxel size was 0.2 x 0.2 x 0.45 µm^3^ to satisfy Nyquist sampling requirements. Even illumination of the entire microcarrier surface was achieved using 5 – 15% power with dual-sided objective illumination with pivot scanning and online maximum intensity fusion. Z-stacks were acquired at 50 frames per second (FPS) using pco.edge 5.5M sCMOS cameras.

### Light-sheet tomography

Light-sheet tomography based on elastic scattering contrast was performed on the Z.1 LSM with the same illumination and detection optics. The laser blocking filter and emission filters were removed from the optical path between the detection objective lens and the camera to allow scattered light to enter the detection path. Samples were illuminated using the 638 nm laser at 0.1% power which was sufficient to fill the dynamic range of the camera at 50 FPS. Imaging was performed with dual-sided objective illumination with pivot scanning and online maximum intensity fusion.

### Mie scattering simulations

The open-source Mie scattering software MiePlot (v4.6.21) was used to simulate the angular distribution of Mie scattering intensity [53]. The software accepts refractive index (n) inputs for the immersion medium and particle, particle size, illumination wavelength, and illumination polarization. The polystyrene particles were modeled as a sphere with 160 µm diameter and n equal to 1.59 [54], and the gelMA microcarriers were spheres with 120 µm diameter and n equal to 1.35 [47]. Cells were simulated to have a diameter of 10 µm and n equal to 1.45 [55], [56]. The combined water and agarose immersion media was modeled with n equal to 1.34. The refractive indices of the agarose immersion media and gelMA microcarriers were measured using a handheld refractometer (KRUSS Optronic).

### Data analysis

The open-source image analysis software ImageJ/FIJI was used to generate 2D intensity projections of the 3D microcarrier fluorescence and scattering volumes. ImageJ was also used to characterize fluorescence signal attenuation with increasing depths by creating an orthogonal projection of the microcarriers and extracting intensity profiles at varying depths of the microcarrier. MATLAB 2020b was used for statistical analysis of the normalized maximum intensity depth profiles of the two microcarriers and the simulated 100% confluent microcarrier. The optical surface area per slice of a sphere was calculated by integrating the sphere surface area using the following equation:

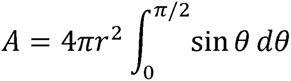

## Results

### Mie scattering simulations

To assess the potential for elastic scattering-based contrast imaging of cells expanded on spherical microcarriers, Mie simulations for the two microcarriers were performed using MiePlot. The angular distributions of Mie scattering intensity for the two microcarriers and a cell were simulated using an unpolarized light source (Figure 1a). Additionally, the gelMA microcarrier was simulated as 160 μm diameter to match the diameter of and compare to the polystyrene microcarrier (Figure 1b).

**Figure 1.**
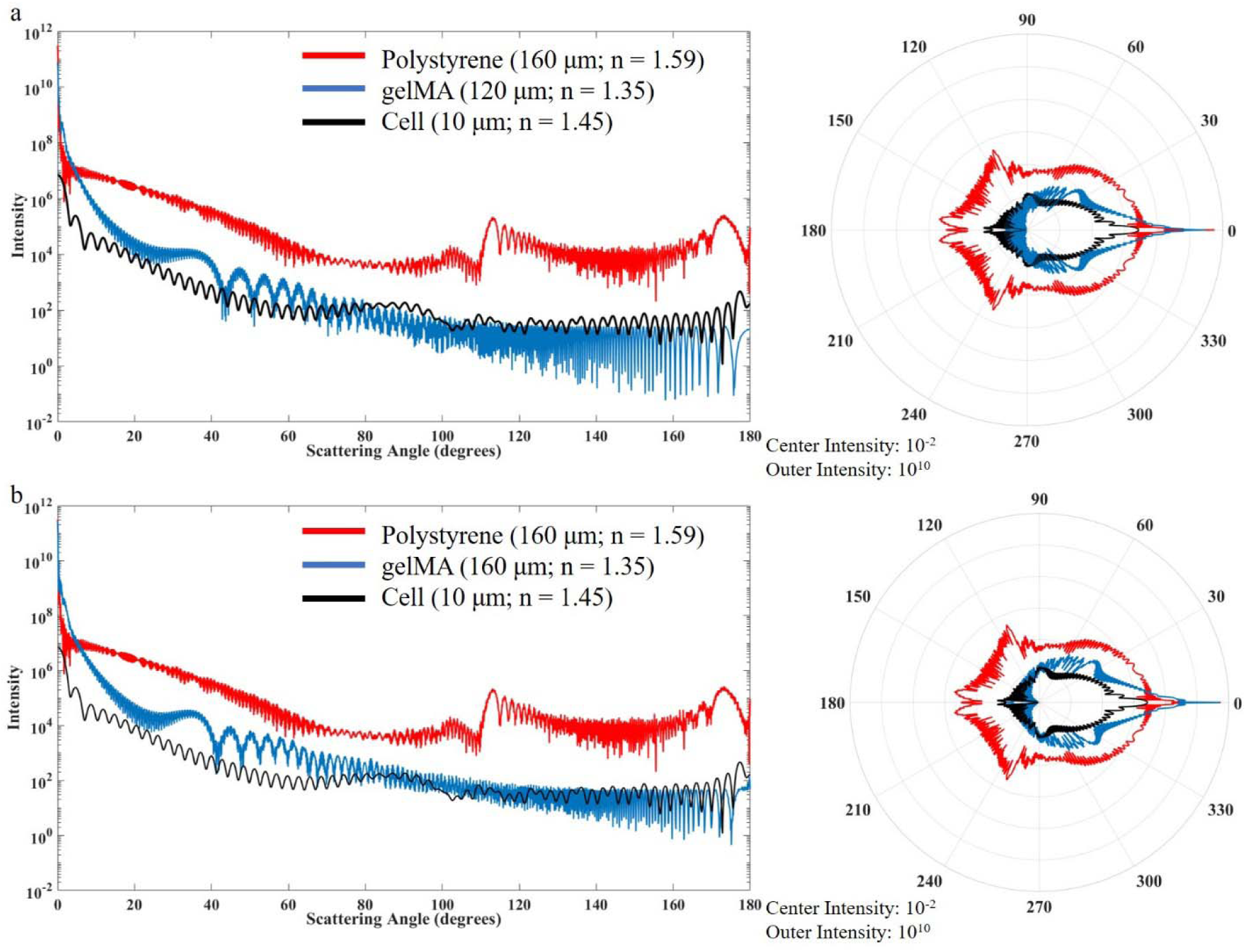
Mie scattering intensity simulations of polystyrene and gelMA microcarriers and single cells on a) rectangular and polar plot showing the polystyrene microcarrier orthogonal scatters at much higher intensities which inhibits visualization of cells along the surface using LSM. b) The gelMA microcarrier was simulated as the same diameter (160 μm) as the polystyrene microcarrier to decouple the scattering intensity from microcarrier size.

These simulations show that the polystyrene microcarriers scatter light much more intensely than the hydrogel microcarriers and the cells. This phenomenon is due to the refractive index mismatch of the surrounding water immersion media and the microcarrier. The Mie simulations predict the scattering intensity to be 2-4 orders of magnitude greater for polystyrene than gelMA microcarriers. The cells, with an overall refractive index between water and polystyrene, scatter light near the same intensity range as the hydrogel microcarriers. This phenomenon permits visualization of cells along the entire gelMA microcarrier surface using both fluorescence and elastic scattering contrast.

### Microcarrier screening for label-free elastic scattering-based visualization

The polystyrene microcarrier scatters much more intensely than cells cultured on the microcarrier surface. This causes the cell signal to be obscured by the polystyrene microcarrier scattering (Figure 2a). Only cells situated on the polystyrene microcarrier equator and proximal hemisphere can be visualized using brightfield transmitted illumination (Figure 2b). This necessitates the use of fluorescence contrast to visualize and distinguish cells from the polystyrene microcarrier surface (Figure 2c). The gelMA microcarrier, conversely, scatters at intensities near or lower than cells cultured on the gelMA microcarrier surface which permits sub-cellular visualization of cells along the entire 3D surface using light-sheet tomography (Figure 2d). Standard deviation and sum of slices contrast enhancement projections even permit simultaneous visualization of the microcarrier, cells, and surrounding agarose. Brightfield illumination can also be used to visualize and identify cells from the gelMA microcarrier surface (Figure 2e). Populated microcarriers can be distinguished from empty microcarriers using brightfield illumination to visualize textural features on the microcarrier surface. Fluorescence contrast similarly provides an equivalent view of cells cultured on the gelMA microcarrier surface (Figure 2f). These exemplary volumetric projections illustrate the capabilities of the hydrogel microcarrier for label-free visualization of cells along the entire microcarrier surface.

**Figure 2.**
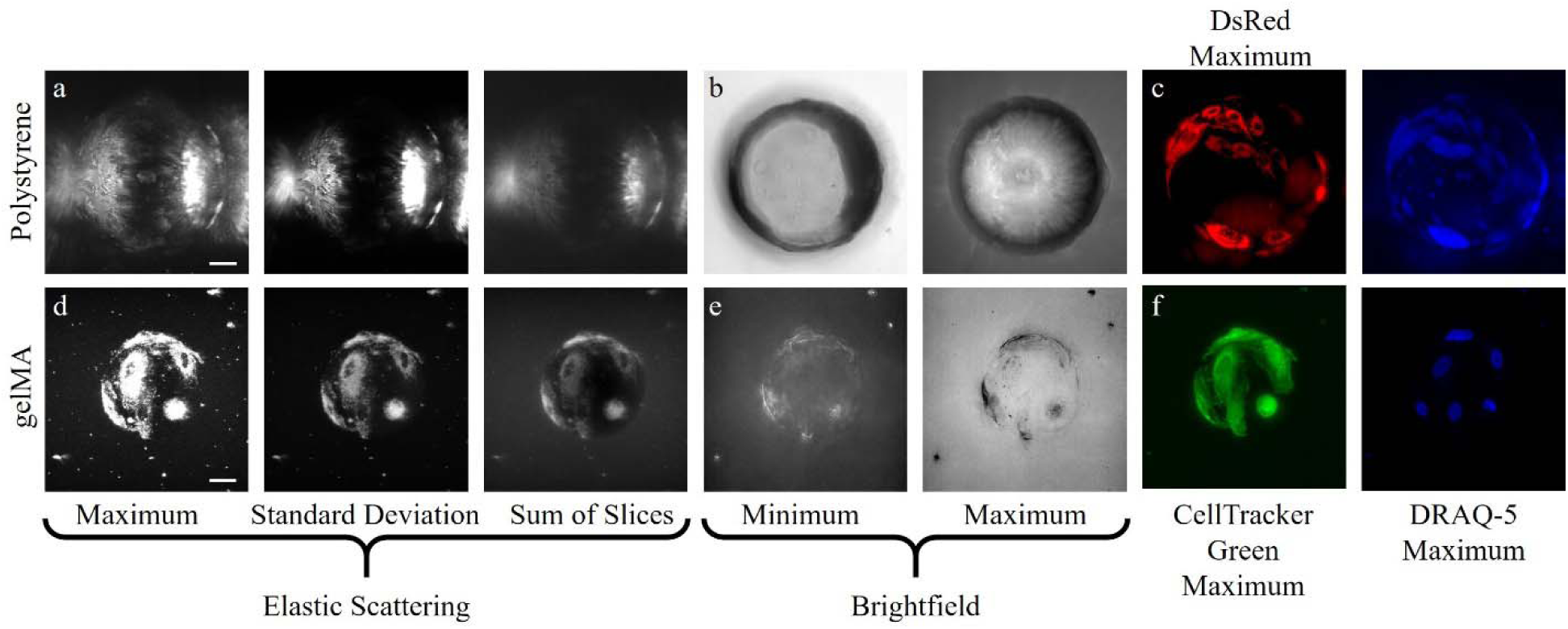
The gelMA microcarriers are superior for label-free visualization of the entire culture surface area. Label-free intensity-based projections of the polystyrene microcarrier using a) elastic scattering and b) brightfield contrast provide limited visualization of cells on the microcarrier surface compared to c) exogenous fluorescence contrast. The gelMA microcarrier permits the use of label-free d) elastic scattering and e) brightfield, and f) exogenous fluorescence contrast for visualization of cells along the entire microcarrier surface.

### Microcarrier screening for direct fluorescence-based cell enumeration

To evaluate the optical imaging performance of the different microcarriers, fixed MSCs attached to commercially available 165 µm diameter polystyrene and 120 µm diameter custom-fabricated hydrogel microcarriers were imaged with a fluorescence light-sheet microscope. Representative data of DsRed and DRAQ-5 labeled BM-hMSCs on polystyrene and CTG and DRAQ-5 labeled ihMSCs on gelMA microcarriers are shown (Figures 3 and 4).

**Figure 3.**
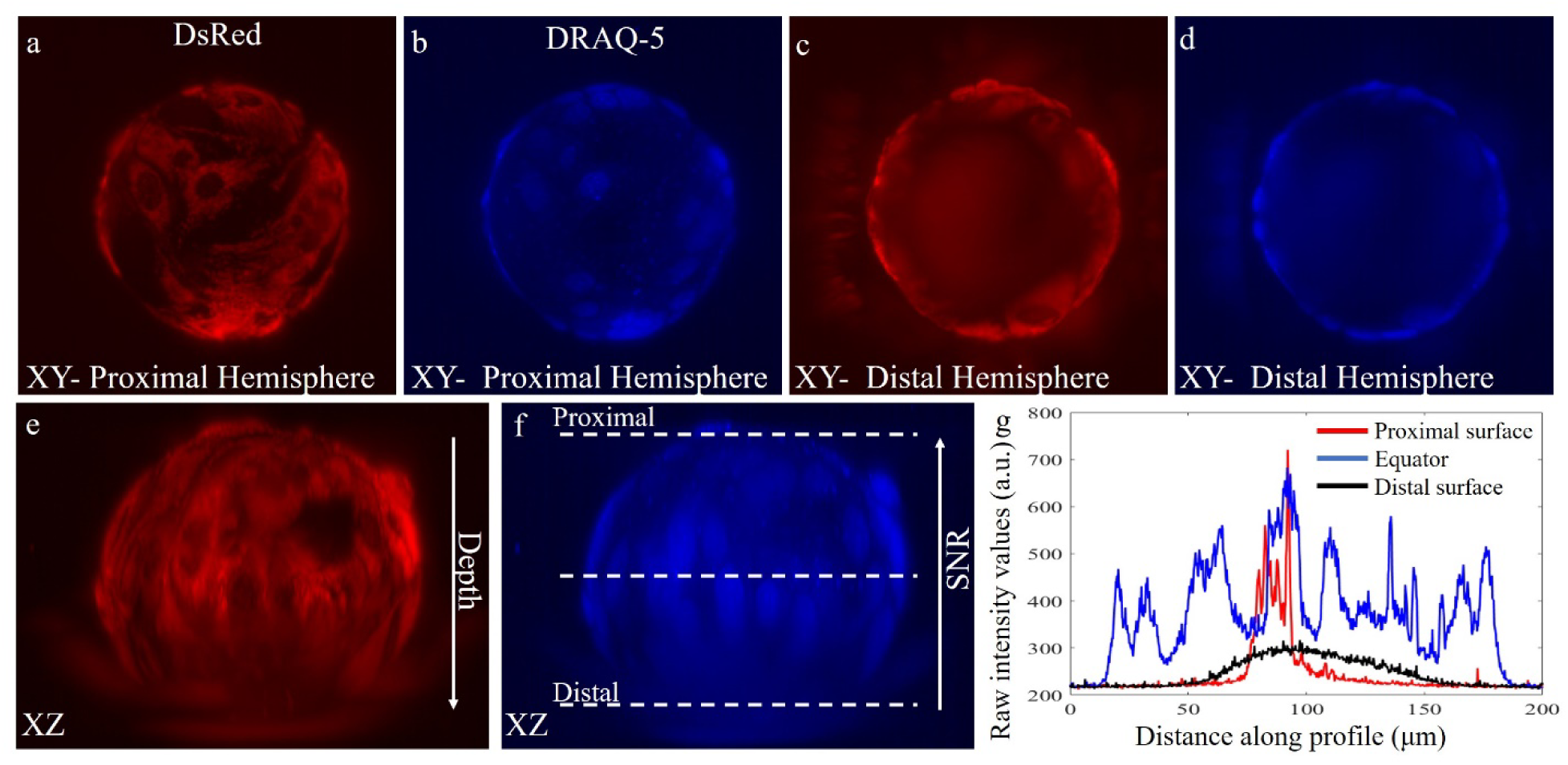
Polystyrene microcarriers limit visualization to the proximal microcarrier surface. Maximum a) dsRed and b) DRAQ-5 intensity projections of the proximal microcarrier surface. c) dsRed and d) DRAQ-5 maximum intensity projections of the distal microcarrier surface. e) dsRed and f) DRAQ-5 signal intensity is g) attenuated beyond the microcarrier equator. Line projections of fluorescence intensity in g correspond to dashed lines in f.

**Figure 4.**
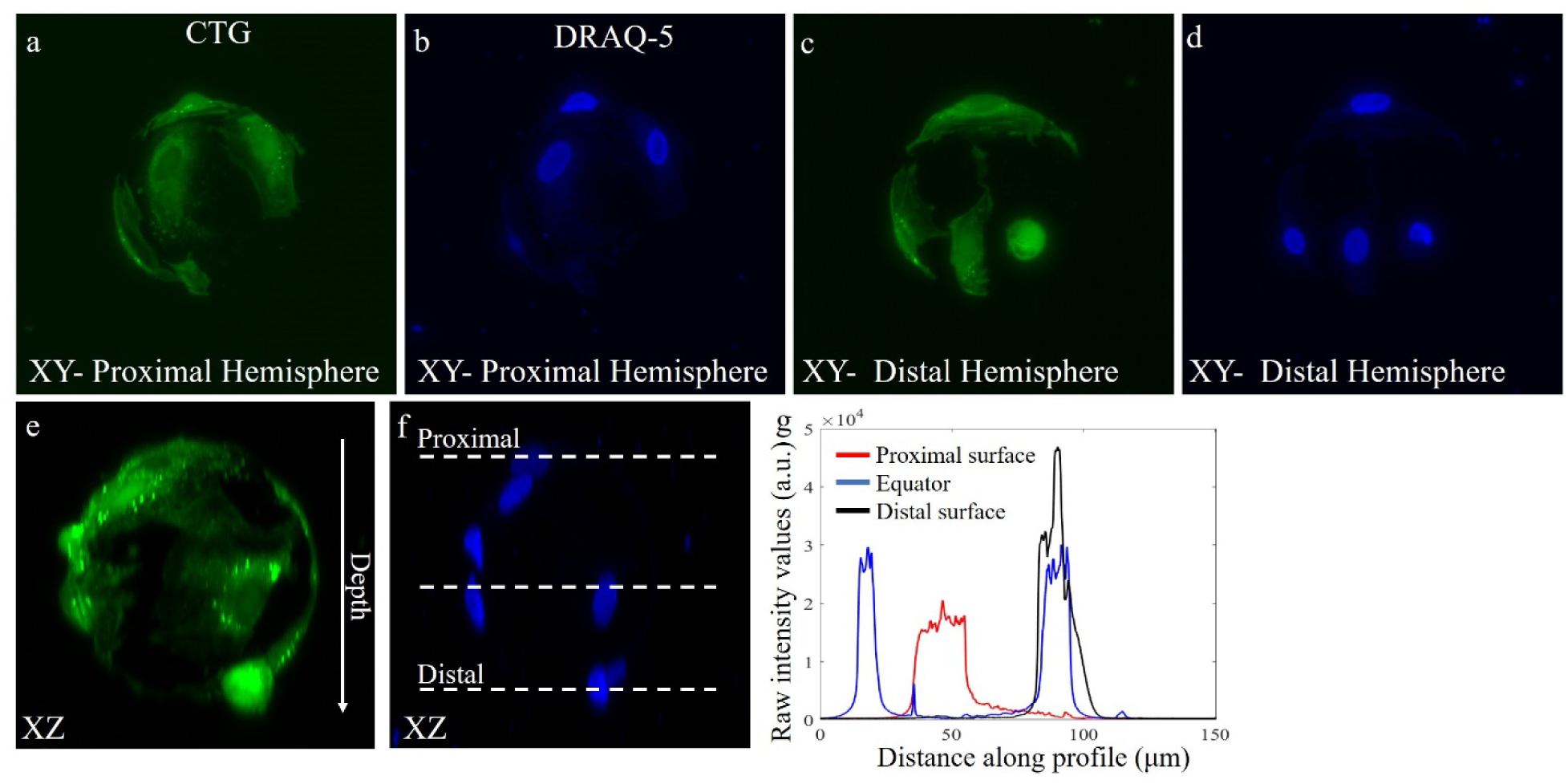
gelMA microcarriers enable high resolution visualization of the entire microcarrier surface. Maximum a) CTG and b) DRAQ-5 intensity projections of the proximal microcarrier surface. c) CTG and d) DRAQ-5 maximum intensity projections of the distal microcarrier surface. e) CTG and f) DRAQ-5 signal intensity is maintained throughout the entire depth of the microcarrier. Line projections of fluorescence intensity in g correspond to dashed lines in f.

The polystyrene microcarrier inhibits visualization of cells on the distal half of the microcarrier equator (Figure 3a-f). There are 30 cell nuclei identified via DRAQ-5 on the microcarrier equator and proximal surface, and only 1 or 2 additional cells can be identified on the distal surface near the equator and no cell nuclei are resolved in the center of the distal microcarrier hemisphere (Figure 3b, 3d). This is because the fluorescence signal intensity is attenuated with depth, and the ability to visualize, resolve, and enumerate cells beyond the microcarrier equator is severely hindered (Figure 3e-f). Cells on the distal half of the microcarrier are illuminated, but the polystyrene microcarrier blurs these slices in the stack and the ability to resolve individual nuclei is eliminated (Figure 3g). A scattering artifact caused by the microcarrier can be seen on the left hemisphere of the microcarrier in Fig. 3c and 3d.

The hydrogel-based gelMA microcarriers, compared to commonly used polystyrene microcarriers, enable high fidelity visualization and semi-automatic direct enumeration of cells along the entire microcarrier surface (Figure 4a-d). There are 7 cell nuclei resolved from the DRAQ-5 contrast (Figure 4f). The fluorescence intensity is preserved along the entire depth of the microcarrier, and cell nuclei can be visualized on both the proximal and distal hemisphere of the microcarrier (Figure 4e-g).

### Depth-dependent fluorescence signal attenuation

To compare the depth-dependent fluorescence signal intensity attenuation caused by the microcarrier material, the maximum DRAQ-5 intensity per frame per microcarrier profile is averaged, normalized, and plotted for both microcarrier materials; additionally, the theoretical illuminated microcarrier surface area is plotted as a reference for a 100% confluent microcarrier. Only single microcarriers with two or more cells and >20% CTG confluency were included to select for moderate and highly confluent single microcarriers and to avoid including aggregated microcarriers in the analysis. The gelMA microcarriers have a more symmetrical profile than the polystyrene microcarriers, and these data show that the gelMA microcarriers cause less depth-dependent fluorescence signal intensity attenuation than the polystyrene microcarriers (Figure 5). A two-sample Kolmogorov-Smirnov test was performed to quantitively assess the similarity of the depth-intensity profiles between the polystyrene and gelMA microcarriers. The gelMA and polystyrene microcarriers showed a statistically significant difference from each other (*P value* = 0.0314). While the gelMA microcarrier profile did not show statistically significant difference from the 100% confluent microcarrier (*P value =* 0.0994), the difference between the polystyrene and 100% confluent microcarrier profiles was found to be statistically significant (*P value =* 0.0082). Data were considered to be significant if the P values were less than 0.05. Both microcarrier material intensity profiles show a decrease in intensity and deviation from the theoretical confluent microcarrier beginning at around 55% depth; this is an interesting phenomenon that may be due the spherical nature of the microcarriers or a sample size artifact. Mie scattering of spherical objects is well-known to have both broad and sharp intensity oscillations termed wiggles and ripples, respectively, that are due to the spherical shape of the scattering object [57], [58]; the depth-dependent fluorescence intensity oscillations seen here could similarly be related to the spherical nature of the microcarriers.

**Figure 5.**
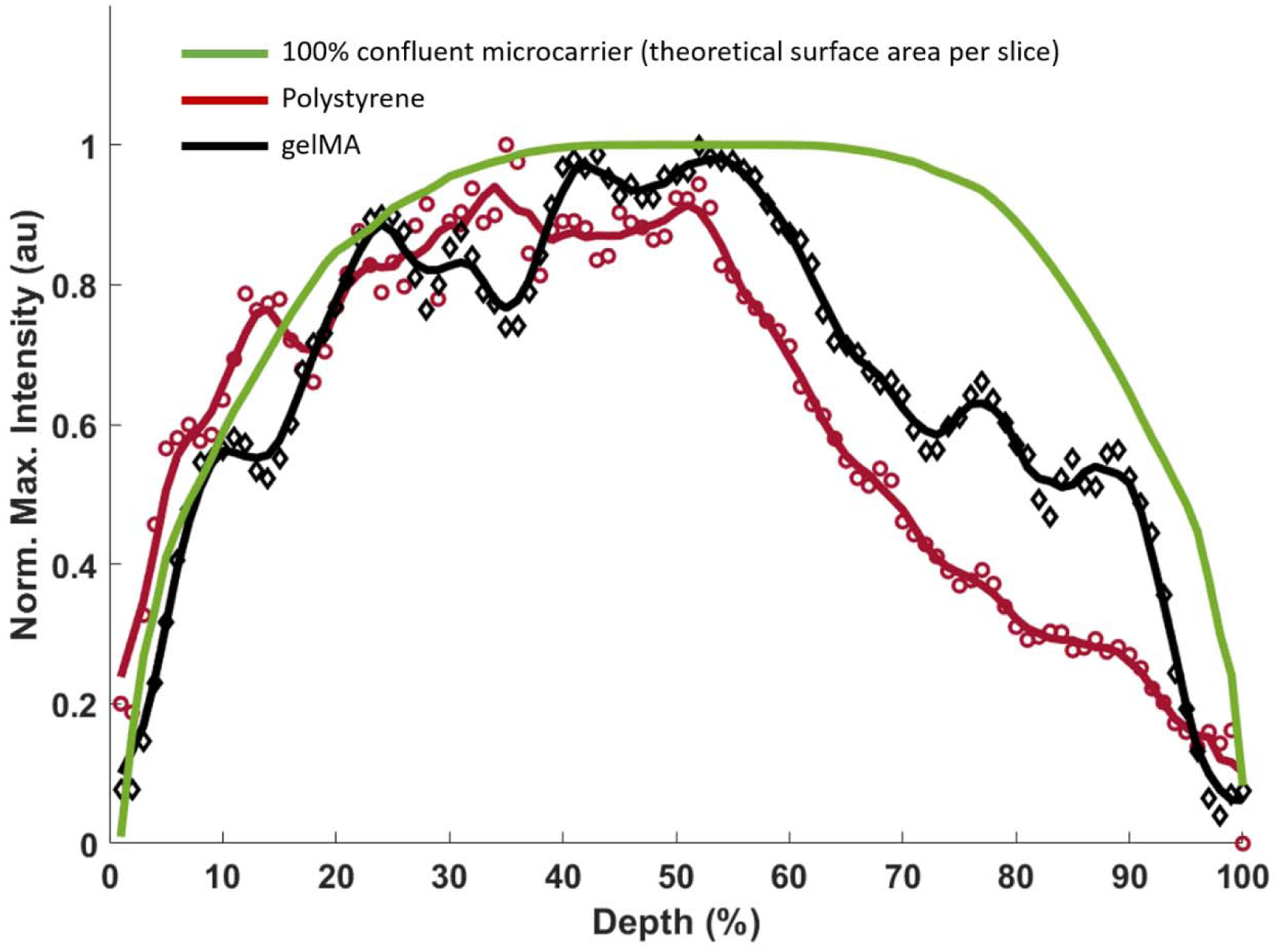
gelMA microcarriers have superior volumetric optical imaging capabilities compared polystyrene microcarriers. Normalized average maximum intensity-depth profile of gelMA (n=17) and polystyrene (n=31) microcarriers showing the large fluorescence intensity attenuation that occurs on the distal half of the polystyrene microcarriers compared to gelMA microcarriers. A 5-point sliding average filter was applied to both microcarrier profiles (solid lines). A 100% confluent microcarrier is simulated using the theoretical surface area per optical slice.

## Discussion

The FDA has encouraged therapeutic manufacturers to deploy PATs as a means to monitor and control a manufacturing process by gathering (near) real-time insight into critical attributes of the therapeutic product that can be incorporated into a control strategy to assure product quality [59]. The combination of optically-compatible microcarriers, rapid, label-free, high-resolution imaging, and robust image analysis-based characterization of adherent cell cultures will greatly enhance the ability to monitor large-scale expansion of cells for cell-based therapies. In this study, we compared the optical imaging capabilities of commercial polystyrene microcarriers versus custom-fabricated gelatin methacrylate (gelMA) microcarriers for non-destructive and non-invasive visualization of the entire microcarrier surface, direct cell enumeration, and sub-cellular visualization of MSCs using Mie scattering simulations and empirical light-sheet imaging data. The Mie scattering simulations explain previous results using reflectance confocal microscopy (RCM) and light-sheet tomography for scattering-based volumetric imaging of microcarrier cell cultures [46], [47]; the polystyrene microcarrier scattering intensity obscures the cell scattering signal and fluorescence modalities are unable to visualize the distal hemisphere of the polystyrene microcarrier. Image formation from the distal microcarrier hemisphere requires photons scattered by cells to reach the camera, but these signal-contributing photons are likely to be deflected and/or absorbed by the polystyrene microcarrier before they reach the detector. Light-sheet imaging is traditionally performed with de-coupled and orthogonal illumination and detection paths, and the detection objective used here has a 97.5° angular aperture. It is apparent from the Mie scattering plots that the polystyrene microcarrier scatters more intensely than the gelMA microcarrier in the detection objective lens angular aperture range, as well as in the forward direction which is relevant for transmitted illumination imaging schemes.

The imaging data illustrate the limited optical imaging capabilities of commonly employed polystyrene microcarriers for *in toto* quantitative monitoring of ihMSC growth on microcarriers. The polystyrene microcarrier does not permit visualization of the entire 3D surface due to its refractive index mismatch, high scattering, and absorption of illumination light, and thus observation of cells is limited to proximal hemisphere of the spherical microcarrier. The custom gelMA microcarriers are optically transparent with a refractive index close to water which allows for label-free visualization of the entire microcarrier core and surface. These experimental findings are supported by the Mie scattering simulations, which showed the value in refractive index matching of the microcarrier for label-free scattering-based 3D imaging of MSCs. Additionally, cell nuclei fluorescently labeled with DRAQ-5 situated anywhere on the microcarrier surface can be visualized with high spatial resolution and SNR as there is minimal refractive index mismatch between the gelMA microcarrier material and the surrounding agarose and DI water suspension that would cause optical aberrations. Cell enumeration is, therefore, no longer limited to the superficial half of the microcarrier surface nor does it require the membrane lysis or detachment of cells from the microcarrier surface.

Volumetric optical imaging enables high fidelity visualization of the entire microcarrier surface. Coupled with hydrogel microcarriers, cells can be visualized using fluorescence and elastic scattering contrast. The use of light-sheet tomography and gelMA microcarriers permits faster acquisition and visualization of adherent cells using multiple orders of magnitude less power than is required for fluorescence contrast. The angular scattering intensity distribution profiles of the microcarriers were simulated to assess the potential for elastic scattering-based imaging of the cells cultured on the microcarrier surface. Additionally, the optical imaging capabilities of polystyrene and gelMA microcarriers were quantitatively compared and show a significant difference in the ability to visualize and characterize the entire microcarrier surface.

## Conclusion

Optically transparent microcarriers composed of gelMA, or commercially available Cytodex 3 composed of dextran, for example [60], [61], enable non-invasive and non-destructive *in toto* quantification of cell density and microcarrier surface area confluency [43], [45]. These optical-quality microcarriers permit light-sheet fluorescence imaging of the entire microcarrier surface. The polystyrene microcarrier causes greater scattering and absorption of illumination light than hydrogel microcarriers, and this prevents visualization of cells on the distal half of the microcarrier using either fluorescence or elastic scattering contrast. Fluorescent labeling of cells renders them unsuitable for additional culture and passaging in a cytotherapeutic manufacturing process. Therefore, label-free contrast is essential for in- or on-line monitoring of cell cultures to not only preserve the sterility of the culture and minimize waste of valuable resources and product, but also for rapid results to enable real-time decision making in the upstream manufacturing process. The combination of optical-quality gelMA microcarriers and label-free light-sheet tomography will facilitate enhanced control of the bioreactor-microcarrier cell culture processes and aid in the development of PATs for cytotherapy manufacturing quality control.

